# A Legionella effector kinase is activated by host inositol hexakisphosphate

**DOI:** 10.1101/2020.02.18.954925

**Authors:** Anju Sreelatha, Christine Nolan, Brenden C. Park, Krzysztof Pawłowski, Diana R. Tomchick, Vincent S. Tagliabracci

## Abstract

The transfer of a phosphate from ATP to a protein substrate, a modification known as phosphorylation, is catalyzed by protein kinases. Protein kinases play a crucial role in virtually every cellular activity. Recent studies of atypical protein kinases have highlighted the structural similarity of the kinase superfamily despite notable differences in primary amino acid sequence. We searched for putative protein kinases in the intracellular bacterial pathogen, *Legionella pneumophila* and identified the Type-4 secretion system (T4SS) effector, Lpg2603 as a remote member of the protein kinase superfamily. We show that Lpg2603 is an active protein kinase with several atypical structural features. Importantly, we find that the eukaryotic-specific host signaling molecule, inositol hexakisphosphate (IP6) is required for Lpg2603 kinase activity. Crystal structures of Lpg2603 in the apo-form and bound to IP6 reveal active site rearrangement that allows for ATP binding and catalysis. Our results on the structure and activity of Lpg2603 reveal a unique mode of regulation of protein kinases and will aid future work into the function of this effector during *Legionella* pathogenesis.

## Introduction

Protein kinases are a class of enzymes that catalyze phosphorylation, a post translational modification (PTM) involving the transfer of the terminal phosphate from ATP to protein substrates (1). The human genome encodes over 500 kinases that play a crucial role in cellular function and are implicated in virtually every cellular activity (2). The protein kinase superfamily can be broadly divided into 2 groups: the eukaryotic protein kinases and the atypical protein kinases. Eukaryotic protein kinases consist of an N-lobe and a C-lobe that harbor conserved amino acid motifs necessary for catalytic function. In contrast, atypical protein kinases lack easily detectable sequence similarity to the eukaryotic protein kinases but still retain catalytic activity. Several atypical members of the kinase superfamily have emerged in recent years that share the structural kinase fold despite significant sequence divergence (3).

We have taken a bioinformatics approach to analyze and identify these atypical and uncharacterized members of the protein kinase superfamily. With this strategy, we identified the Golgi casein kinase, Fam20C, that phosphorylates serine residues within the Ser-x-Glu/pSer consensus motif, found in roughly 75% of human plasma and cerebrospinal fluid phosphoproteins (4,5). Using sequence similarity to Fam20C, we identified the atypical protein kinase, CotH, present in many bacterial and eukaryotic spore forming organisms. CotH phosphorylates spore coat proteins for effective spore germination in *Bacillus subtilis* (6). Furthermore, we identified a distant member of the kinase superfamily, the SelO family, that is conserved from bacteria to humans and catalyzes protein AMPylation instead of phosphorylation (7).

A subset of proteins that our bioinformatics approach continues to bring to the forefront are bacterial effector proteins. Bacterial effector proteins serve as a diverse pool of proteins in which many atypical and novel biological mechanisms can be found (8). HopBF1, for example, is a family of bacterial effector proteins from *Pseudomonas syringae* that utilize a novel molecular ‘mimicry’ to phosphorylate host cell Hsp90 to evade the host immune defense during infection (9). Likewise, *Legionella pneumophila*, a gram-negative intracellular pathogen known to be the causative agent of Legionnaires disease, contains a plethora of effector proteins with exciting biology. *L. pneumophila* secretes over 300 effectors to alter the host cellular processes to form a replicative niche and evade degradation (10,11). One such effector protein, SidJ, also retains a protein kinase-like fold but catalyzes protein polyglutamylation that is dependent on the host cofactor calmodulin (12-15).

Interestingly, the *L. pneumophila* genome encodes for 5 eukaryotic-like protein kinases that manipulate host cell signaling (16,17). LegK1-4 and LegK7 are serine/threonine kinases that target various host pathways including actin remodeling, protein synthesis, the immune response, the protein folding machinery and the Hippo pathway (18-23). In search of atypical protein kinases in *L. pneumophila*, we identified a kinase domain in the T4SS effector protein, Lpg2603 (lem28, sdmB) that has also been predicted to have a kinase fold by Burstein et al. (24). Previously, a conserved phosphatidylinositol-4-phophate (PI4P) binding domain at the C terminus of Lpg2603 was identified and shown to localize the protein to the *Legionella* containing vacuole (LCV) during infection (25). This localization is shared by *L. pneumophila* T4SS effectors, Lpg1101/Lem4 and DrrA/sidM, which harbor the conserved PI4P binding domain at their C-termini but differ in their N-terminal domains (25). DrrA is a multidomain effector with a nucleotidyltransferase domain and a guanine nucleotide exchange factor (GEF) domain in addition to the PI4P binding domain (26,27). In contrast, Lpg1101 contains an N-terminal haloacid dehalogenase (HAD) domain with tyrosine phosphatase activity (28). However, the structure and biochemical activity of Lpg2603 remained uncharacterized.

Here, we show that Lpg2603 is an active protein kinase with several unusual structural features not typical of canonical protein kinases. Importantly, we found that Lpg2603 requires binding of the host cofactor inositol hexakisphosphate (IP6) for activation through a unique mode of active site rearrangement. Our results not only provide insights into the mechanism by which Lpg2603 catalyzes phosphorylation, but also highlight a unique mechanism of allosteric regulation of kinases by IP6.

## Results

### Lpg2603 has sequence similarity to protein kinases

A bioinformatic screen for *Legionella* effectors bearing sequence similarity to protein kinases was performed using the FFAS algorithm (29) and yielded Lpg2603 as a likely protein kinase, with 10-18% sequence identity to known bacterial kinases ospG, NleH, YopO, ppkA and several human kinases. The FFAS alignments allowed unequivocal assignments of the typical features of protein kinases: the Gly-rich ATP-binding loop as well as the predicted catalytic Asp201 and Mg^2+^ ion-binding Asp225 (**Fig. 1*A***). In contrast, the ion pair Lys72 and Glu91 (Protein Kinase A nomenclature) are absent in Lpg2603 (**Fig. S1**). Homologs of Lpg2603 are found only in a range of strains from the *Legionella/Fluoribacter* genus.

**Figure 1.**
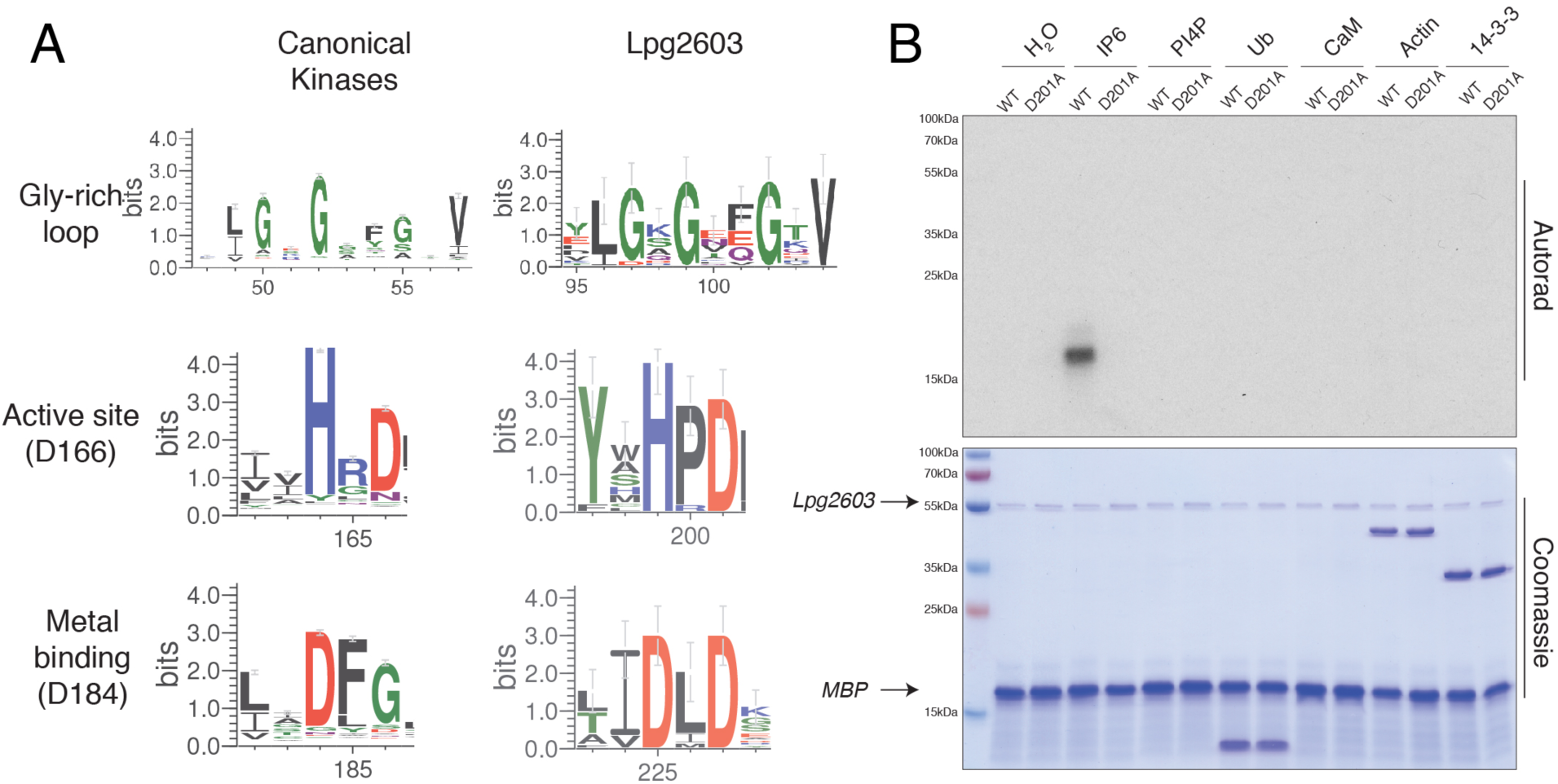
Lpg2603 is an active kinase in the presence of inositol hexakisphosphate (IP6). **(A)** Sequence logo built using 74 homologs of Lpg2603 depicting conserved motifs present in kinases including the Gly-rich loop and active site residues (numbering in parentheses corresponds to PKA). **(B)** Autoradiograph depicting the incorporation of ^32^P from [γ-^32^P] ATP into myelin basic protein (MBP) by recombinant Lpg2603 wild type (WT) or the inactive mutant, D201A, in the presence of different cofactors: inositol hexakisphosphate (IP6), phosphatidylinositol 4-phosphate (PI4P), ubiquitin (Ub), calmodulin (CaM), globular actin or 14-3-3 protein. Reaction products were resolved by SDS-PAGE and visualized by Coomassie blue staining (lower) and autoradiography (upper).

### Lpg2603 is an active kinase

To determine whether Lpg2603 is an active kinase, we expressed *L. pneumophila* Lpg2603 in *Escherichia coli* as a 6X-His-Sumo-fusion protein and purified the protein by Ni-NTA affinity chromatography. We also purified recombinant Lpg2603 containing an alanine mutation in the predicted catalytic Asp201 (PKA nomenclature Asp166). Following removal of the 6X-His-Sumo tag, we performed kinase assays in the presence of [γ^32^P]ATP. Recombinant *L. pneumophila* Lpg2603, however, did not phosphorylate the generic protein kinase substrate, myelin basic protein (MBP) (**Lanes 1 and 2, Fig. 1*B***). Bacterial effectors often utilize eukaryotic specific cofactors to regulate their activity within the host cell while remaining inactive in the bacterial cell (30). Therefore, we tested the activity of Lpg2603 in the presence of some common cofactors. Remarkably, wildtype (WT) Lpg2603, but not the predicted catalytically inactive D201A mutant, phosphorylated MBP in the presence of inositol hexakisphosphate (IP6) but none of the other cofactors that we tested (**Fig. 1*B***).

### Lpg2603 requires IP6 for optimal activity

Given the structural similarity in inositol phosphates, we tested if other inositol phosphates could activate Lpg2603. Lpg2603 displayed a strong preference for IP6 in comparison to myo-inositol, IP, IP2, IP3, IP4 and IP5 with optimal MBP phosphorylation observed at approximately 65 μM IP6 (**Fig. 2*A* and 2*B***). To determine the binding affinity of IP6 to Lpg2603 we measured the dissociation constant (Kd) using isothermal titration calorimetry (ITC). IP6 bound to Lpg2603 with a Kd of ∼315 μM, which is in accordance with eukaryotic cellular IP6 concentration of up to 1 mM (**Fig. 2*C***) (31).

**Figure 2.**
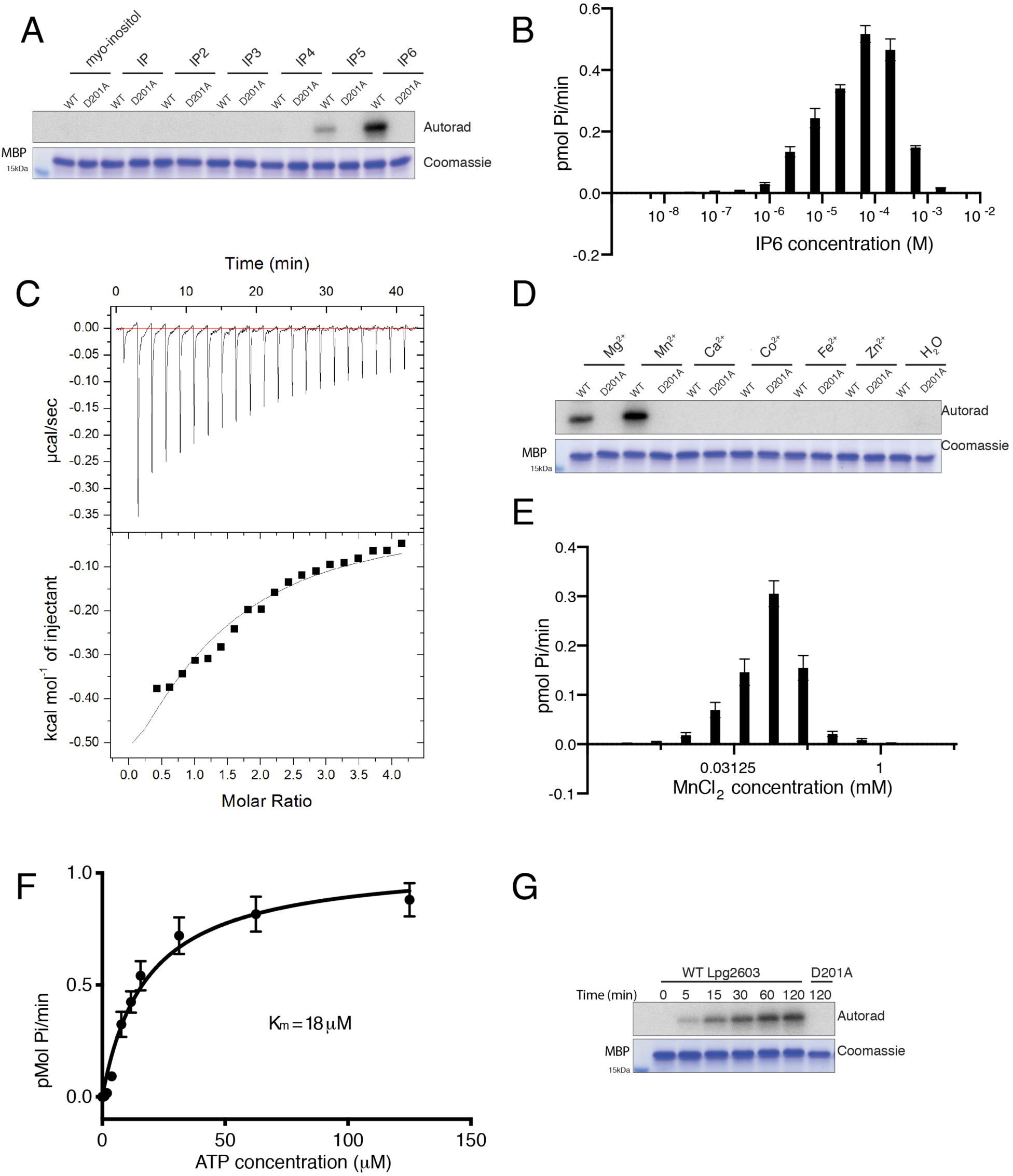
Lpg2603 activity requires IP6 and Mn^2+^. **(A)** *In vitro* kinase assay showing incorporation of ^32^P from [γ-^32^P] ATP into MBP by Lpg2603 in the presence of myo-inositol, inositol phosphate (IP), inositol diphosphate (IP2), inositol triphosphophate (IP3), inositol tetraphosphate (IP4), inositol pentaphosphate (IP5), inositol hexakisphosphate (IP6). Reaction products were analyzed as in **Figure 1B**. **(B)** Incorporation of γ-^32^P from [γ-^32^P] ATP into MBP by Lpg2603 with increasing concentrations of IP6. Reaction products were resolved by SDS-PAGE and Coomassie stained MBP bands were excised for scintillation counting. **(C)** Representative isothermal calorimetry data for Lpg2603 binding to IP6. Lpg2603 is at 200 μM in the cell and IP6 is at 8mM in the titration syringe for a final molar ratio of 1:4. **(D)** *In vitro* kinase assay showing incorporation of γ-^32^P from [γ-^32^P] ATP into MBP by Lpg2603 in the presence or absence of MgCl_2_, MnCl_2_, CaCl_2_, CoCl_2_, FeCl_2_, ZnCl_2_. Reaction products were analyzed as in **Figure 1B**. **(E)** Incorporation of γ-^32^P from [γ-^32^P] ATP into MBP by Lpg2603 with varying concentrations of MnCl_2_. Reaction products were analyzed as in **Figure 2B**. **(F)** Kinetic analysis depicting the concentration dependence of Mn^2+^/ATP on the rate of MBP phosphorylation by Lpg2603. Reaction products were analyzed as in **Figure 2B**. *K*_*m*_ for Mn^2+^/ATP = 18.09µM; *Vmax* = 1.053pmol/min. **(G)** Time-dependent incorporation of γ-^32^P from [γ-^32^P] ATP into MBP by Lpg2603. Reaction products were analyzed as in **Figure 1B.**

We next assayed the optimal divalent metal for kinase activity. Lpg2603, but not the inactive mutant, phosphorylated MBP in the presence of Mg^2+^and Mn^2+^ (**Fig. 2*D***). Lpg2603 demonstrates a preference for Mn^2+^ which, interestingly, is inhibitory at high concentrations (**Fig. 2*E***). Similar to canonical kinases, Lpg2603 has a K_m_ for ATP of 18μM (**Fig. 2*F***). Furthermore, Lpg2603, but not the inactive mutant, phosphorylated MBP in a time-dependent manner (**Fig. 2*G***). Collectively, our results demonstrate that Lpg2603 is a bacterial effector kinase that requires IP6 for catalytic activity.

### Crystal structure of Lpg2603 reveals IP6 binding pocket and mechanism of activation

To gain further insight into the mechanism of Lpg2603 activity, we solved the crystal structure of a fragment of Lpg2603 which lacks the PI4P binding region at a resolution of 2.10 Å (**Fig. 3*A***). The apo structure displayed a unique N-terminal extension that is rich in β-strands. There was no observable electron density for residues 76-117 of the N-lobe indicating a flexible protein region. Electron density is absent for residues that are important for canonical kinase activity including the Gly-rich loop and ion pair Lys from the VAIK motif. The C-lobe consists of the five α-helices including the catalytic Asp201 and metal binding Asp225. The apo structure can be superimposed onto PKA with an RMSD of 4.8 Å over 178 C-α backbone atoms.

**Figure 3.**
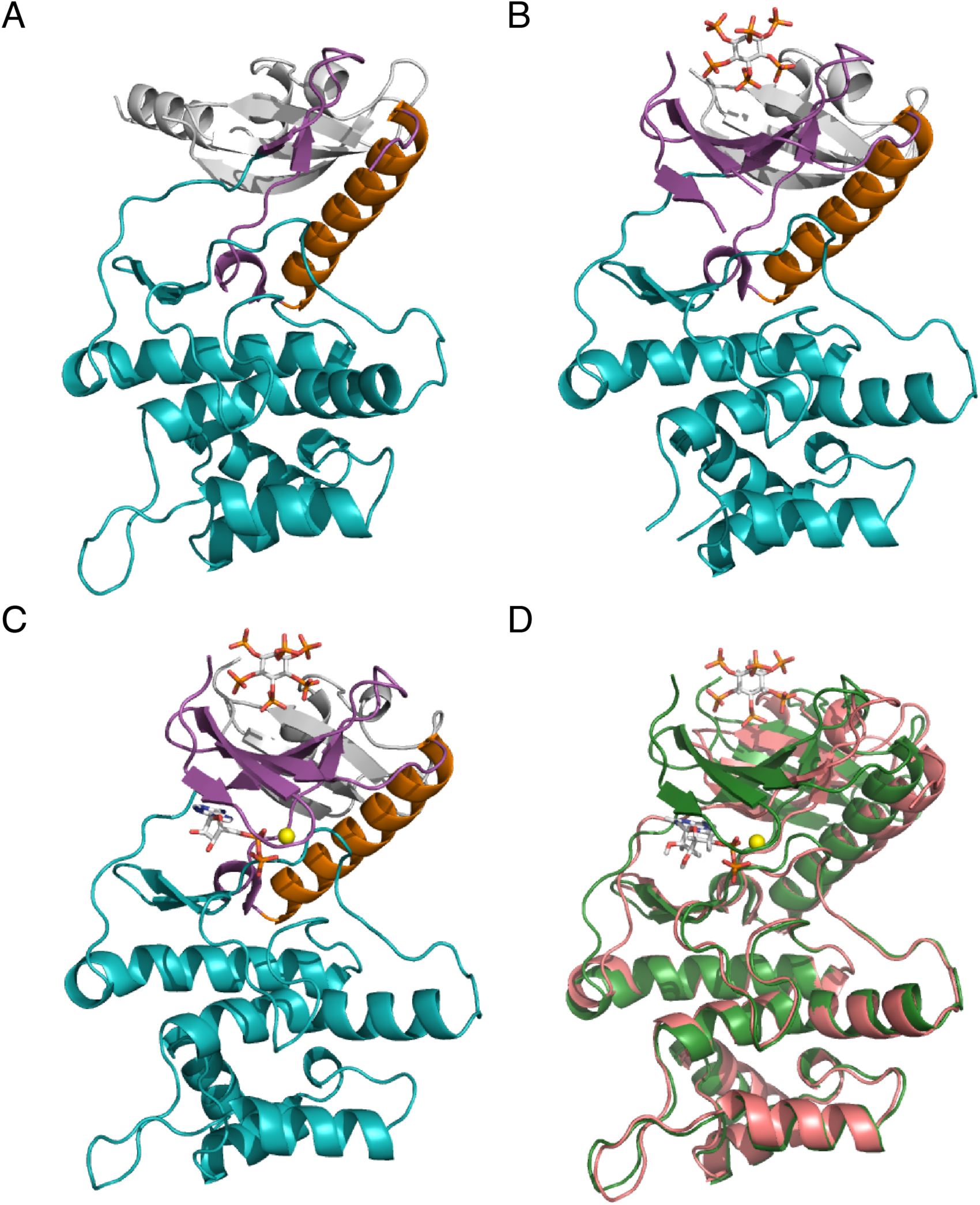
Crystal structure of Lpg2603 reveals a unique mode of kinase activation by IP6. **(A)** Ribbon representation of apo-Lpg2603. The N-lobe and C-lobe are depicted in magenta and teal, respectively. The αC-helix and the N-terminal extensions are shown in orange and white, respectively. **(B)** Ribbon representation of Lpg2603 bound to IP6. The ligand IP6 is shown in ball-and stick, colored according to atom. **(C)** Ribbon representation of Lpg2603 bound to IP6 and ADP. The nucleotide ADP and ligand IP6 are shown in ball-and stick, colored according to atom. The Mn^2+^ ion is shown as a yellow sphere. **(D)** Superposition of the apo-Lpg2603 (shown in pink) with the nucleotide and ligand bound Lpg2603 (shown in green) depicting the disordered N-lobe in the apo structure.

Next, we sought to compare the apo structure to the IP6 bound structure to determine the effect of cofactor binding. Surprisingly, the crystal structure of Lpg2603 bound to IP6 revealed a kinase fold with a highly ordered N-lobe with clear electron density for residues 92-117, which form three of the five strands of the canonical N-lobe β-sheet (**Fig. 3*B***). The IP6 is coordinated in a positively charged pocket composed of residues from the N-lobe β-sheet (corresponding to strands β1 to β5 of classical kinases) and residues from the N-terminal extension that includes a three-stranded β-sheet. The IP6 binding site is very well conserved in Lpg2603 homologs suggesting a common mechanism of activation (**Fig. S1**). While the C-lobes of the apo and IP6 bound structures remain similar, conformational shifts are observed in the IP6-binding cradle, αC-helix, and metal binding residues that favor ATP binding (**Fig. 3*A* and *B***). The apo structure can be superimposed onto the IP6 bound structure with an RMSD of 2.3 Å over 251 C-α backbone atoms. Thus, we hypothesize that binding of IP6 stabilizes the N-lobe of Lpg2603 to allow for ATP binding and kinase activity.

We next solved the crystal structure of Lpg2603 bound to IP6 and ADP to identify the residues that facilitate nucleotide coordination upon IP6 binding (**Fig. 3*C***). The overall conformation of the kinase in the presence and absence of nucleotide is very similar with an RMSD of 0.7 Å over 274 C-α backbone atoms (**Fig. 3*D***). One notable difference upon ADP binding is the ordering of the Gly-rich loop that folds over the nucleotide in the active site (**Fig. S2**). Structural homology searches using DALI identified membrane-associated tyrosine-and threonine-specific cdc2-inhibitory kinase (Myt1), troponin I-interacting kinase (TNNI3K), and Bruton’s tyrosine kinase (BTK) as the closest structural homologs of Lpg2603.

### Structure guided mutagenesis highlight the residues important for nucleotide and IP6 coordination

Using the Lpg2603 holoenzyme structure, we identified key residues that form the IP6 binding pocket (**Fig. 4*A***). In order to identify IP6 binding residues that are important for kinase activity, we purified recombinant Lpg2603 proteins with alanine substitutions at Lys114, Lys156, Asn154, Lys107, Arg50, and Lys76. As expected, mutations in these residues to an Ala either weakened or completely abolished Lpg2603 activation (**Fig. 4*B***). These results support our hypothesis that proper coordination of IP6 is necessary for Lpg2603 activation.

**Figure 4.**
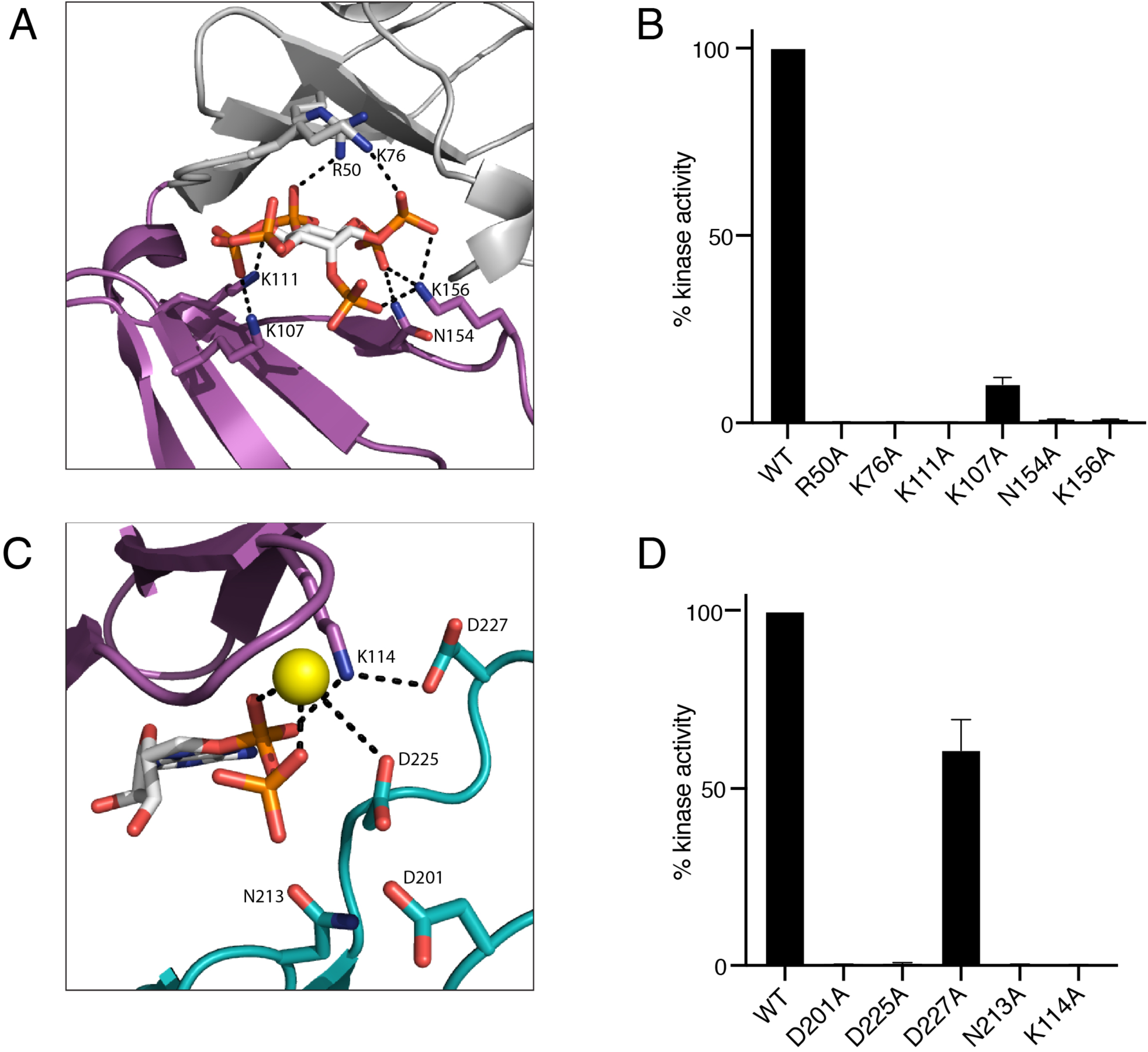
Mutational analysis highlights residues in Lpg2603 important for catalysis. **(A)** Enlarged image of IP6-binding pocket showing residues and interactions important for IP6 binding. IP6 is shown in ball-and stick, colored according to atom. **(B)** Activity of Lpg2603 or mutants were assayed as in **Figure 2B**. Activity is expressed relative to the WT enzyme. **(C)** Enlarged image of the nucleotide-binding pocket showing residues and interactions important for ADP binding and catalysis. The bound ADP is shown in ball-and stick, colored according to atom. Mn^2+^ ion is shown as a yellow sphere. **(D)** Lpg2603 or mutants were assayed as in **Figure 2B**. Activity is expressed relative to the WT enzyme.

Next, we assayed recombinant Lpg2603 proteins with mutations in the nucleotide binding pocket (**Fig. 4*C***). The highly conserved catalytic loop in Lpg2603 consists of a non-canonical HPD motif with Asp201 likely acting as the catalytic base. Mutation of the catalytic Asp201 abolished kinase activity of Lpg2603 (**Fig. 4*D***). A metal binding DxD motif is present in Lpg2603 where the Asp225 coordinates the Mn^2+^ ions in the active site. Mutation of the predicted metal binding residues, Asp225 and Asn213 eliminated activity of the kinase. Lys114 from the β3 strand (PKA equivalent K72) coordinates the α and β phosphates of ATP (**Fig. 4*C***). However, the glutamate corresponding to the canonical Glu91(PKA) from the αC-helix that forms the conserved K72-E91 ion pair is positioned away from the active site while Asp227 coordinates the ion pair Lys114. Interestingly, mutation of Lys114 completely abolished kinase activity while mutation of Asp227, that forms an ion pair with Lys114, retained more than 50% of activity. This is reminiscent of the hypomorphic E74A ion pair mutant of the bacterial effector kinase HopBF1 (9). Collectively, our results highlight the residues important for IP6 binding and phosphotransfer.

### IP6 binding facilitates nucleotide binding in Lpg2603

Given the structural reorganization observed with IP6, we investigated whether IP6 binding is required for nucleotide binding. In order to assay for ATP binding, we performed ITC with the weakly hydrolysable ATP analog, AMP-PNP. In the absence of IP6, Lpg2603 no longer binds to the nucleotide (**Fig. 5*A***). Upon addition of IP6, Lpg2603 binds to AMP-PNP with a Kd of 3.6 μM (**Fig. 5*B***). The IP6 bound Lpg2603 crystal structure and the *in vitro* kinase assays indicate Lys111 as a key residue for IP6-induced activation of kinase activity. Lpg2603 K111A does not bind to AMP-PNP in the presence or absence of IP6 (**Fig. 5*C* and *D***). These results provide further evidence that IP6 allosterically activates Lpg2603 kinase activity.

**Figure 5.**
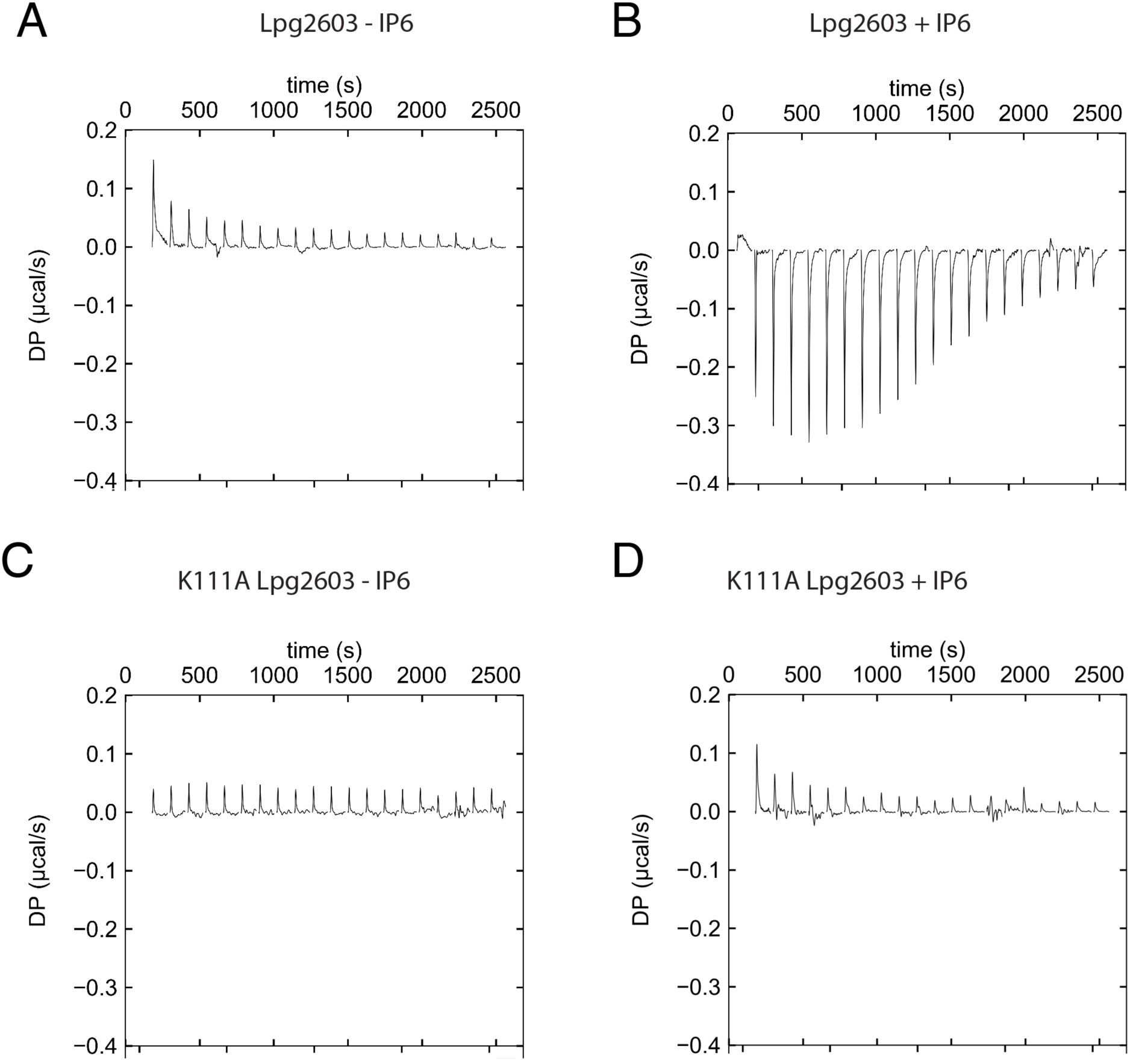
Nucleotide binding to Lpg2603 is dependent on IP6 binding. **(A)** Representative isothermal titration calorimetry data for Lpg2603 binding to AMP-PNP in the absence of IP6. For all instances, Lpg2603 is at 100uM in the cell and AMP-PNP is present at 2mM in the titration syringe to a final molar ratio of 1:4. **(B)** Representative isothermal titration calorimetry data for Lpg2603 binding to AMP-PNP in the presence of IP6. **(C)** Representative isothermal titration calorimetry data for Lpg2603 K111A binding to AMP-PNP. **(D)** Representative isothermal titration calorimetry data for Lpg2603 K111A binding to AMP-PNP in the presence of IP6.

## Discussion

Lpg2603 harbors a conserved *Legionella*-effector PI4P binding region at the C-terminus, which is required for anchoring the kinase to the *Legionella* containing vacuole during an infection (25). We have modeled this domain by using the homologous PI4P binding domain from the *Legionella* effector DrrA and docked it against the kinase domain (**Fig. S3**). The full-length structure model displays the lipid binding domain adjacent to the C-lobe of the kinase, suggesting that PI4P binding and/or membrane localization may affect kinase function of Lpg2603. This also suggests that its substrates may be membrane anchored or membrane-proximal. Despite our efforts to identify interacting proteins, we were unable to determine the host substrate. Nevertheless, our results of the IP6-dependent kinase activation will pave the way to identify the *in vivo* substrate during *Legionella* infection.

We have identified a new mechanism of bacterial effector kinase regulation by a host signaling molecule, IP6. IP6 is a eukaryote-specific ligand that is absent in bacteria (32). Interestingly, it is also the most abundant inositol phosphate in eukaryotic cells where it regulates several processes including growth factor signaling, cell cycle progression, and vesicle trafficking (31). Cleverly, bacterial effectors have evolved to hijack host signaling molecules such as IP6 to spatially regulate activity (30). Less than a handful of other bacterial effectors from different pathogens are activated by IP6 including acetyl transferases (HopZ1,YopJ and AvrA) and a cysteine protease (VPA1380) (30). Thus, the bacterial enzyme remains inactive within the bacterial cell until it is delivered to its destination within the eukaryotic host cell. Notably, this is the first example of a bacterial effector kinase that requires IP6 for activation. The IP6 binding site of Lpg2603 is spatially distinct from the kinase active site suggesting allosteric regulation rather than IP6 acting as a cofactor for catalysis (**Fig. 3*C***). Hence, we hypothesize that binding of IP6 at the N-terminal extension triggers conformational changes in multiple amino acids which functionally link the IP6 binding pocket to the kinase active site.

In addition to bacterial effectors, two eukaryotic protein kinases, Bruton’s tyrosine kinase (BTK) and Casein kinase 2 (CK2), have also been shown to be activated by IP6 binding (33,34). BTK is a non-receptor tyrosine kinase that undergoes IP6-induced dimerization and activation (34). BTK is critical for proper B cell function and mutations are implicated in chronic lymphocytic leukemia, X-linked agammaglobulinemia, and several other autoimmune diseases (35). BTK is composed of multiple domains: Pleckstrin Homology-Tec Homology (PH-TH) domain, kinase domain, SH2 and SH3 domain (36). IP6 binds to the PH-TH domain to induce dimerization and trans-autophosphorylation of the kinase domains, leading to activation (34). Notably, the mechanism of IP6 activation of BTK is distinct from Lpg2603 as we did not observe any dimerization of Lpg2603 upon incubation with IP6 (data not shown). In contrast to BTK, IP6 binds to the basic patch in the substrate recognition site of the C-lobe in CK2 (33). CK2 is a serine/threonine kinase that regulates cell proliferation (37); however, the physiological relevance of regulation of CK2 by IP6 is unclear.

In comparison with the IP6 binding site of BTK (4Y94) and CK2 (3W8L), the IP6 binding site in Lpg2603 is more tightly coordinated by charged residues (**Fig. 4*A* and 6**). Moreover, in BTK and CK2, the sites are rather shallow and are built by residues from two different monomers (**Fig. 6**). The IP6 binding pocket in Lpg2603 consists of amino acids from the N-terminal extension and the N-lobe of the kinase that are highly conserved within homologs of Lpg2603. Furthermore, the apo structure revealed a fairly disordered N-lobe and active site in the absence of IP6, highlighting the importance of binding this ligand for activity. Hence, our crystallographic and biochemical analysis reveals a previously undocumented mode of allosteric regulation of the kinase superfamily.

**Figure 6.**
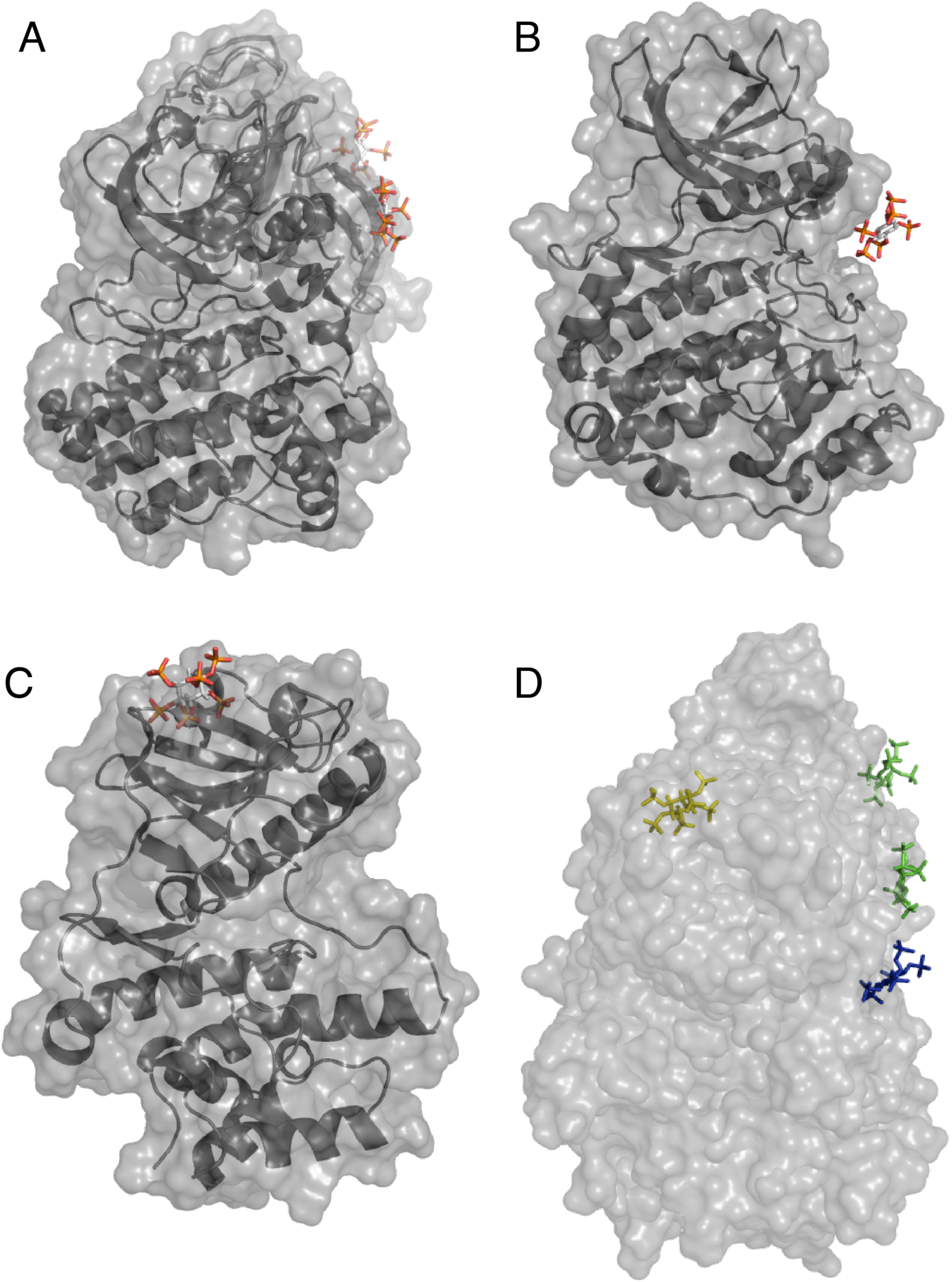
Structural comparison reveals unique mode of IP6 binding to Lpg2603. **(A)** Surface representation of BTK bound to IP6. PDB 4Y93,4Y94. **(B)** Surface representation of CK2 bound to IP6. PDB 3W8L. **(C)** Surface representation of Lpg2603 bound to IP6. **(D)** Surface representation of the superposition of BTK, CK2 and Lpg2603. IP6 bound to BTK, CK2 and Lpg2603 is shown in green, blue and yellow, respectively.

### Experimental Procedures Reagents

Selenomethionine media was purchased from Molecular dimensions (MD12-500). Inositol 1,3,4,5 tetraphosphate (Q-1345), Inositol 1,4 bisphosphate (Q-0014), Inositol 1,4,5 phosphate (Q-0145), phosphatidylinositol 4-phosphate (P4008a) were purchased from Echelon Biosciences. D-myo-inositol-1,3,4,5,6 pentaphosphate (10009851) was purchased from Cayman chemicals. Myo-inositol 1-dihydrogen phosphate (S860360), myo-inositol (I7508), inositol hexakisphosphate (P8810), Myelin basic protein (M1891), and AMP-PNP (A2647) were purchased from Sigma. Ubiquitin and calmodulin were expressed as 6X His fusion proteins and purified from *E. coli* as described (12). g-actin was a generous gift from Michael Rosen (38).

### Generation of Constructs

*L. pneumophila* Lpg2603 residues 10-C, 21-322 or 10-322 Lpg2603 were cloned into a modified pet28a bacterial expression vector (ppSumo), containing an N-terminal 6X-His tag followed by the yeast Sumo (smt3). The coding sequence for the yeast homologue of 14-3-3, BMH2, was amplified by PCR using *Saccharomyces cerevisiae* BY4741 gDNA as a template. The amplified open reading frames were cloned into pGEX4T containing an N-terminal GST tag.

### Protein Expression and Purification

*L. pneumophila* Lpg2603 10-C, 21-322, or 10-322 ppSumo were transformed into *Escherichia coli* Rosetta (DE3) competent cells. Cells were grown in Luria-Bertani (LB) medium at 37°C until the A_600_ reached ∼0.5-0.7. Protein expression was induced by 0.4 mM isopropyl β-D-thiogalactoside (IPTG) overnight at room temperature. The cells were harvested by centrifugation, lysed in 50 mM Tris-HCl pH 8.0, 300 mM NaCl, 1 mM PMSF, and 0.1% β-mercaptoethanol) by sonication. Cell lysates were centrifuged at 25,000 x g for 25 min. The cleared lysate was incubated with Ni-NTAagarose for approximately one hour at 4°C. Beads were passed over a column and washed with 20 column volumes of 50 mM Tris pH 8, 300 mM NaCl, 10 mM imidazole. Protein was eluted off the beads with 50mM Tris pH 8, 300 mM NaCl, 300 mM imidazole. Proteins were cut overnight at 4°C with 6X-His tagged ULP Sumo protease and further purified using Superdex200 size exclusion chromatography column attached to an AKTA Pure FPLC system (GE Healthcare).

For purification of the yeast homolog of 14-3-3, BMH2 pGEX4T was transformed into *Escherichia coli* Rosetta (DE3) competent cells. Cells were grown in Luria-Bertani (LB) medium at 37°C until the A_600_ reached ∼0.5-0.7. Protein expression was induced by 0.4 mM isopropyl β-D-thiogalactoside (IPTG) overnight at room temperature. The cells were harvested by centrifugation, lysed in 50 mM Tris-HCl pH 8.0, 300 mM NaCl, 1 mM PMSF, and 0.1% β-mercaptoethanol) by sonication. Cell lysates were centrifuged at 25,000 x g for 25 min. The cleared lysate was incubated with glutathione sepharose for approximately one hour at 4°C. Beads were passed over a column and washed with 20 column volumes of 50 mM Tris pH 8, 300 mM NaCl, 0.1% β-mercaptoethanol. Protein was eluted off the beads with 50mM Tris pH 8, 300 mM NaCl, 10 mM reduced glutathione.

### In vitro kinase assays

*In vitro* kinase assays were performed using untagged Lpg2603 10-C in a reaction mixture containing 50 mM Tris pH 7.5, 100 µM MnCl_2,_ 100 µM [γ-^32^P] ATP (SA = 1000 cpm/pmol), 1 mM DTT, 167 µg/mL MBP, 21 µg/mL Lpg2603 and 20 µM IP6. Reactions were incubated at 25°C for 10 minutes and terminated by the addition of 0.5 mM EDTA. SDS loading buffer was added to the samples and boiled. Reaction products were separated by SDS-PAGE and visualized by Coomassie blue staining. For comparison of cofactors, reactions were performed as above with the following modifications: reactions contained 20 µM IP6, 188 µM PI4P, 40 µg/mL ubiquitin, 40 µg/mL calmodulin, or 40 µg/mL BMH2.

For comparison of inositols, reactions were performed as above with the following modifications: reactions were incubated with 20 µM of the indicated inositols.

For comparison of metals, reactions were performed as above with the following modifications: reactions contained 100 µM of the indicated metals.

For comparison of Lpg2603 point mutants, reactions were performed as above with the following modifications: reactions contained 100 µM [γ-^32^P] ATP (SA = 5000 cpm/pmol).

For the kinetic analysis, reactions were performed as above with the following modifications: reactions contained 300 µM MnCl_2_, 300 µM [γ-^32^P] ATP (SA = 5000 cpm/pmol). Reactions were incubated at 25°C for 15 minutes.

### Crystallization and structure determination

For apo-Lpg2603, recombinant Lpg2603 10-322 in 5 mM Tris-HCl pH 8, 30 mM NaCl was concentrated to 10 mg/mL. The crystals were grown at 20°C by the sitting drop vapor diffusion method using a 1:1 ratio of protein:reservoir solution containing 18% PEG3350, 0.2 M LiCl, and were flash frozen in 20% PEG3350, 0.2 M LiCl, 30 mM NaCl and 30% ethylene glycol. Apo-Lpg2603 crystals exhibited the symmetry of space group P3_1_21 with cell dimensions of a = 62.96 Å, c = 176.41 Å, contained one apo-Lpg2603 per asymmetric unit, and diffracted to a minimum Bragg spacing (d_min_) of 2.10 Å when exposed to synchrotron radiation. Selenomethionine labeled protein was obtained by expressing Lpg2603 10-322 in B834 cells grown in SelenoMet™ media. SeMet Lpg2603 10-322 in 5 mM Tris-HCl pH 8, 30 mM NaCl was concentrated to 20 mg/mL. The crystals were grown at 20°C by the sitting drop vapor diffusion method using 1:1 ratio of protein:reservoir solution containing 13% PEG3350, 0.2 M LiCl, and were flash frozen in 15.5% PEG3350, 0.2 M LiCl, 30 mM NaCl and 35% ethylene glycol. SeMet apo-Lpg2603 crystals exhibited the symmetry of space group P3_1_21 with cell dimensions of a = 63.11 Å, c = 175.51 Å, contained one SeMet apo-Lpg2603 per asymmetric unit, and diffracted to a minimum Bragg spacing (d_min_) of 2.15 Å when exposed to synchrotron radiation.

For IP6 bound Lpg2603, Lpg2603 10-322 in 5 mM Tris-HCl pH 8, 30 mM NaCl, and 1mM IP6 was concentrated to 16 mg/mL. The crystals were grown at 20°C by the sitting drop vapor diffusion method using a 1:1 ratio of protein:reservoir solution containing 0.1 M sodium acetate pH 5.75, and were flash frozen in 0.1 M sodium acetate pH 5.75, 30 mM NaCl, 1mM IP6 and 45% ethylene glycol. IP6 bound Lpg2603 crystals exhibited the symmetry of space group P2_1_2_1_2_1_ with cell dimensions of a = 52.93 Å, b = 73.21 Å, c = 261.25 Å, contained three IP6 bound Lpg2603 molecules per asymmetric unit, and diffracted to a minimum Bragg spacing (d_min_) of 2.65 Å when exposed to synchrotron radiation.

For IP6 and ADP bound Lpg2603, SeMet Lpg2603 21-322 in 5 mM Tris-HCl pH 8, 30 mM NaCl, 1 mM DTT, 0.5 mM IP6, 1 mM MnCl_2_, 1 mM AMP-PNP was concentrated to 10 mg/mL. The crystals were grown at 20°C by the sitting drop vapor diffusion method using a 1:1 ratio of protein:reservoir solution containing 0.1 M citric acid pH 4.0, 6% MPD. Wells were allowed to equilibrate for approximately 24 hours and crystal growth was initiated by micro-seeding. Crystals were flash frozen in 0.1 M citric acid pH 3.5, 7% MPD, 30 mM NaCl, 0.5 mM IP6, 1 mM MnCl_2_, 1 mM AMP-PNP, and 35% ethylene glycol. IP6 and ADP bound Lpg2603 crystals exhibited the symmetry of space group P2_1_2_1_2 with cell dimensions of a = 52.54 Å, b = 77.07 Å, c = 72.56 Å, contained one IP6 and ADP bound Lpg2603 per asymmetric unit, and diffracted to a minimum Bragg spacing (d_min_) of 1.77 Å when exposed to CuKα radiation from a home source. Although we added AMP-PNP to the protein, we observed ADP in our crystal structure. This may be due to contaminating ATP or weak hydrolysis of AMP-PNP.

Diffraction data were collected at 100 K at the Advanced Photon Source beamline 19-ID for all datasets except the IP6 and ADP bound Lpg2603, which was collected with CuKa radiation from a Rigaku Xtal MM003 source. Anomalous data for the SeMet apo-Lpg2603 were collected near the Se K-edge. Data were indexed, integrated, and scaled using the *HKL-3000* program package (39). Data collection statistics are provided in **Table S1**.

### Phase determination and structure refinement

Phases for SeMet apo-Lpg2603 were obtained from a single-wavelength anomalous dispersion experiment using a selenomethionyl-derivatized protein crystal with data collected at the selenium K-edge to a d_min_ of 2.15 Å. Seven selenium sites were located using the program *SHELXD* (40), and phases were refined with the program *SHELXE* (41) resulting in an over-all figure-of-merit of 0.66 for data between 46.39 and 2.15 Å. Phases were further improved by density modification in the program dm (42). An initial model containing 74% of all apo-Lpg2603 residues was automatically generated in the program ARP/wARP (43).

As the selenomethionyl derivatized and native crystals were isomorphous, all further calculations for the native structure were performed versus the native data. Additional residues for apo-Lpg2603 were manually modeled in the program Coot (44). Positional and isotropic atomic displacement parameter (ADP) as well as TLS ADP refinement was performed to a resolution of 2.10 Å using the program Phenix (45) with a random 5% of all data set aside for an R_free_ calculation. The current model contains one apo-Lpg2603 monomer; included are residues 1 – 75, 118 – 332 and 196 water molecules. The R_work_ is 0.176 and the R_free_ is 0.211. A Ramachandran plot generated with Molprobity (46) indicated that 97.4% of all protein residues are in the most favored regions and none in disallowed regions.

Phases for the IP6 bound Lpg2603 were obtained by the molecular replacement method in the program Phaser (47) using the coordinates for the apo Lpg2603 monomer with residues 1 – 15 removed. Model building and refinement were performed to a resolution of 2.65 Å using a similar protocol to the apo structure. Three IP6 bound Lpg2603 molecules were located in the asymmetric unit, and the electron density for chain C is substantially weaker than for chains A and B. The R_work_ is 0.253, and the R_free_ is 0.282; the presence of anisotropy in the data and the weak electron density for chain C is likely the cause of the higher than expected R_work_ and R_free_ for this model. A Ramachandran plot generated with Molprobity indicates that 95.4% of all protein residues are in the most favored regions and none in disallowed regions.

Phases for the IP6 and ADP bound Lpg2603 were obtained by the molecular replacement method in the program Phaser using the coordinates for the IP6 bound Lpg2603 monomer. Model building and refinement were performed to a resolution of 1.77 Å using a similar protocol to the apo structure. One IP6 and ADP bound Lpg2603 molecule was located in the asymmetric unit. The R_work_ is 0.213, and the R_free_ is 0.237. A Ramachandran plot generated with Molprobity indicates that 95.4% of all protein residues are in the most favored regions and none in disallowed regions.

Phasing and model refinement statistics for all structures are provided in **Table S1**.

### Lpg2603 Sequence Logo

The similarity of Lpg2603 to kinases was identified by screening the set of *Legionella pneumophila* subsp Philadelphia effectors using the FFAS server (48). Homologs of the Lpg2603 kinase domain were collected using BLAST and aligned by Mafft (49). Sequence logos were produced using the Weblogo 3.0 server (50).

### Structural modeling of full length Lpg2603

The structural model of the DrrA domain was built using the 4mxp structure as template for the I-Tasser server (51). The full length structure model of Lpg2603 was built by docking together the kinase-like domain structure and the DrrA domain structure model using the AIDA server (52).

### Structural comparisons

Structures were visualised and analysed in Pymol. Structure database searches were conducted using the Dali and Fatcat servers (53,54).

### Isothermal Calorimetry

Recombinant Lpg2603 10-C wild type or Lpg2603 10-C K111A was produced as described above in *Protein Expression and Purification*. For AMP-PNP binding assays, the final buffer consisted of 50 mM Tris HCl pH 7.5, 150 mM NaCl, 1 mM DTT, 1 mM MgCl_2_, and 1 mM IP6. For IP6 binding, MgCl_2_ and IP6 were excluded from the buffer.

For AMP-PNP binding assays, 2 mM AMP-PNP was titrated into Lpg2603 10-C at 100 µM cell concentration using Malvern MicroCal iTC200 System at 20°C. Initial injection was 0.5 µL and the following 20 injections were 1.9 µL with stirring at 750 rpm. Initial injections were not included in the final analysis. Spacing between injections were 120 seconds to allow for proper baseline equilibration. Resulting thermograms were integrated using the NITPIC software and SEDPHAT was used to fit the isotherms assuming a binary interaction model.

For IP6 binding assays, 8 mM IP6 was titrated into Lpg2603 10-C at 400 µM cell concentration as above. Initial injections were 0.5 µL, and the following 20 injections were 1.9 µL. Initial injections were, once again, not included in the final analysis. Spacing between injections were adjusted to 300 seconds to allow for proper baseline equilibration. Resulting thermograms were analyzed using the Origin software to obtain Kd and confidence intervals.

## Acknowledgements

We thank Shaeri Mukherjee, Kim Orth, Samantha Yee and members of the Tagliabracci lab for valuable input. We thank Chad Brautigam and Shih-Chia Tso for help with ITC. Results shown in this report are derived from work performed at the Argonne National Laboratory, Structural Biology Center at the Advanced Photon Source. This work was supported by NIH Grants R00DK099254 and (V.S.T.), T32DK007257-37 (A.S.), Welch Foundation Grants I-1911 (V.S.T) and the Polish National Agency for Scientific Exchange scholarship PPN/BEK/2018/1/00431 (K.P.). A.S is the W.W. Caruth, Jr. Scholar in Biomedical Research and a Cancer Prevention Research Institute of Texas Scholar (RR190106). V.S.T. is the Michael L. Rosenberg Scholar in Medical Research, a Cancer Prevention Research Institute of Texas Scholar (RR150033) and a Searle Scholar. Research reported in this publication was supported by the Office of The Director, National Institutes of Health under Award Number S10OD025018. The content is solely the responsibility of the authors and does not necessarily represent the official views of the National Institutes of Health.

The Lpg2603 structures have been deposited in the protein data bank (PDB) with accession codes as follows: apo-Lpg2603: 6VVC; Lpg2603 bound to IP6: 6VVD; Lpg2603 bound to IP6, Mn2+, ADP: 6VVE

## Author contributions

A.S., and V.S.T. designed the experiments. A.S, B.C.P, C.N., D.R.T. and V.S.T conducted the experiments. K.P. performed the bioinformatics. A.S. and V.S.T wrote the manuscript with input from all authors.

